# Nuclear Genome Size is Positively Correlated with Mean LTR Insertion Date in Fern and Lycophyte Genomes

**DOI:** 10.1101/571570

**Authors:** Anthony E. Baniaga, Michael S. Barker

## Abstract

Nuclear genome size is highly variable in vascular plants. The composition of long terminal repeat retrotransposons (LTRs) is a chief mechanism of long term change in the amount of nuclear DNA. Compared to flowering plants, little is known about LTR dynamics in ferns and lycophytes. Drawing upon the availability of recently sequenced fern and lycophyte genomes we investigated these dynamics and placed them in the context of vascular plants. We found that similar to seed plants, mean LTR insertion dates were strongly correlated with haploid nuclear genome size. Fern and lycophyte species with small genomes such as those of the heterosporous *Selaginella* and members of the Salviniaceae had recent mean LTR insertion dates, whereas species with large genomes such as homosporous ferns had old mean LTR insertion dates intermediate between angiosperms and gymnosperms. This pattern holds despite methylation and life history differences in ferns and lycophytes compared to seed plants, and our results are consistent with other patterns of structural variation in fern and lycophyte genomes.

---

Vascular plants exhibit an astonishing diversity of genome sizes relative to other eukaryotes. Plant genomes vary more than 2,400 fold in size, from the minute genomes of *Genlisea tuberosa* [1C=0.06 Gbp (Fleischmann et al. 2014; Rivadavia *et al.* 2013)] to the extremely large genomes of *Paris japonica* [1C=152.2 Gbp (Pellicer *et al.* 2010)]. The rate of plant genome size evolution appears to be lineage specific. For example, the genomes of *Genlisea* spp. range from 0.06-1.7 Gbp, with recent reductions in the smallest genomes following recent whole genome duplication (Fleischmann *et al.* 2014). This stands in contrast to the small genomes of the Selaginellaceae that have a narrower range of genome size from 1C 0.082-0.184 Gbp (Baniaga *et al.* 2016) despite deep divergences among subgenera dating to the Permian-Triassic boundary (Arrigo *et al.* 2013; Kenrick and Crane 1997; Korall and Kenrick 2002; Westrand *et al.* 2016; Zhou *et al.* 2016).

The primary mechanisms driving variation in genome size in vascular plants are whole genome duplication (WGD) and the proliferation of transposable elements. Although WGDs are common and phylogenetically widespread throughout the history of vascular plants (Barker *et al.* 2016; Cui *et al.* 2006; Jiao *et al.* 2011; Kagale *et al.* 2014; Landis *et al.* 2018; Li *et al.* 2015; Li *et al.* 2018), genome size is often reduced following polyploidization by fractionation and diploidization (Freeling *et al.* 2012; Murat *et al.* 2014), partially due to species specific maximum genome size thresholds (Zenil-Ferguson *et al.* 2016). Consequently, most of the variation in genome size is caused by the expansion and contraction of Class I transposable elements, specifically Long Terminal Repeat retrotransposons (LTRs) which increase in copy number via a copy-and-paste mechanism (Feschotte *et al.* 2002). LTRs can comprise a majority of a vascular plant genome (Michael 2014), and total LTR content is positively correlated with genome size in angiosperms (Flavell 1980; Michael 2014; Tiley and Burleigh 2015; Wendel *et al.* 2016). For example, in the extremely small genome of *Utricularia gibba*, LTRs comprise only 2% of the genome (Ibarra-Laclette *et al.* 2013), but in *Zea mays* they comprise more than 75% of the genome (Schnable *et al.* 2009). These dynamics lead LTRs to be the dominant driver of genome size evolution in vascular plants.

Comparative analyses of LTR insertion dates can be used to reveal the dynamics of LTRs and their impacts on genome evolution. It is possible to estimate LTR insertion dates because LTRs have identical 5’ and 3’ ends at initial insertion. After insertion, they evolve neutrally and given an estimate of the synonymous substitution rate for a lineage or taxon the divergence can be used to date LTR insertion (SanMiguel *et al.* 1998). Most angiosperms have young and active LTRs inserted 0-4 mya with few full length LTRs recovered beyond 4 mya in monocots and eudicots (El Baidouri and Panaud 2013). Exceptions to this pattern are found in the genomes of *Fritillaria* spp. These plants have some of the largest known angiosperm genomes, and are characterized by the slow accumulation of diverse repeat elements without their subsequent removal (Kelly *et al.* 2015). Similar LTR age distributions are found in gymnosperms, which also have large genome sizes. Their genomes are characterized by numerous diverse repeats with a uniform distribution of LTR insertion times dating back to more than 60 mya (Kovach *et al.* 2010; Nystedt *et al.* 2013; Voronova *et al.* 2017; Zuccolo *et al.* 2015). The persistence of LTRs over millions of years is a common feature of the largest genomes of seed plants.

In contrast to seed plants, we know relatively little about the dynamics of LTRs and genome size evolution in the ferns and lycophytes. Fern nuclear genome size ranges from 1C=250 Mbp in the heterosporous water fern *Salvinia cucullata* (Li *et al.* 2018) to 1C=147.3 Gbp in the whisk-fern *Tmesipteris obliqua* (Hidalgo *et al.* 2017). On average, most ferns have large genome sizes with a mean of 1C=13.98 Gbp (Clark *et al.* 2016), intermediate between those of gymnosperms and angiosperms. Genome sizes in lycophytes range from the extremely small genomes of 1C=81.5 Mbp in the spike-moss *Selaginella selaginoides* (Baniaga *et al.* 2016) to 1C=11.7 Gbp in the quillwort *Isoetes lacustris* (Hanson and Leitch 2002), with an average of 1.66 Gbp for lycophytes (Leitch and Leitch 2013). Unlike seed plants, nuclear genome size in ferns and lycophytes correlates strongly with chromosome number (Bainard *et al.* 2011; Barker 2013; Clark *et al.* 2016; Leitch and Leitch 2013; Nakazato *et al.* 2008). We may thus expect that LTRs may play a lesser role in the evolution of fern and lycophyte genome size compared to seed plants.

Investigations of LTR dynamics in ferns and lycophytes are in their infancy, but have already revealed some important details. Generally, the proportion of the genome comprised of LTRs is within the range observed in seed plants (Banks *et al.* 2011; Li *et al.* 2018; Van Buren *et al.* 2018; Wolf *et al.* 2015; Xu *et al.* 2018). LTR insertion dates have only been estimated for two heterosporous ferns and species of *Selaginella*. In the heterosporous ferns, which have much smaller genome sizes than their homosporous relatives, a high amount of recent activity is found with relatively low amounts of genetic divergence between LTR 5’ and 3’ ends (Li *et al.* 2018). In *Selaginella* the majority of LTRs are also relatively young (Banks *et al.* 2011; Van Buren *et al.* 2018; Xu *et al.* 2018), and are strongly linked to the high haplotype variation observed in *S. lepidophylla* (Van Buren *et al.* 2018). The role of LTRs in influencing haplotype variation in *Selaginella* may explain the high heterozygosity and assembly difficulties of other *Selaginella* genomes despite their minute nature.

Given that chromosome number is strongly correlated with nuclear genome size in ferns and lycophytes (Bainard *et al.* 2011; Barker 2013; Clark *et al.* 2016; Nakazato *et al.* 2008; Leitch and Leitch 2013), we sought to test if genome size is also well correlated with LTR insertion activity. With the advent of newly available genome-wide data for several fern and lycophyte taxa (Banks *et al.* 2011; Li *et al.* 2018; VanBuren *et al.* 2018; Wolf *et al.* 2015; Xu *et al.* 2018), we were able to explore the LTR dynamics of ferns and lycophytes. The addition of these phylogenetically diverse genomic data allowed us to test if genome size was correlated with mean LTR insertion date in ferns and lycophytes, as demonstrated previously for seed plants (Michael 2014). If genome size is correlated with mean LTR insertion date, then we predict that 1) homosporous ferns should have intermediate LTR insertion dates between angiosperms and gymnosperms, and 2) the heterosporous *Selaginella* with their extremely small genomes and the relatively small genomes of the heterosporous water ferns should have recent LTR insertion dates comparable to angiosperms.

## MATERIALS AND METHODS

### LTR discovery

Genome assemblies used in analyses were downloaded from published and publicly available datasets (Li *et al.* 2018; Phytozome v11; VanBuren *et al.* 2018; Wan *et al.* 2018; Wolf *et al.* 2015; Xu *et al.* 2018). Five homosporous fern taxa (*Ceratopteris richardii, Cystopteris protrusa, Dipteris conjugata, Plagiogyria formosana, Pteridium aquilinum*), two heterosporous fern taxa (*Azolla filiculoides, Salvinia cucullata*), and three lycophyte taxa (*Selaginella lepidophylla, S. moellendorffii, S. tamariscina*) were analyzed. Only *Selaginella* taxa were available to represent lycophytes. In general, the proportion of the genome comprised of LTRs in ferns and lycophytes is similar to that observed in other seed plants (Table 1). In addition to the seven fern taxa and three species of *Selaginella*, five representative angiosperms (*Amborella trichopoda, Sorghum bicolor, Arabidopsis thaliana, Erythranthe guttata, Solanum lycopersicum*) and three representative gymnosperms (*Gnetum montanum, Picea abies, Pinus sylvestris*) were included in our analysis.

**Table 1.**
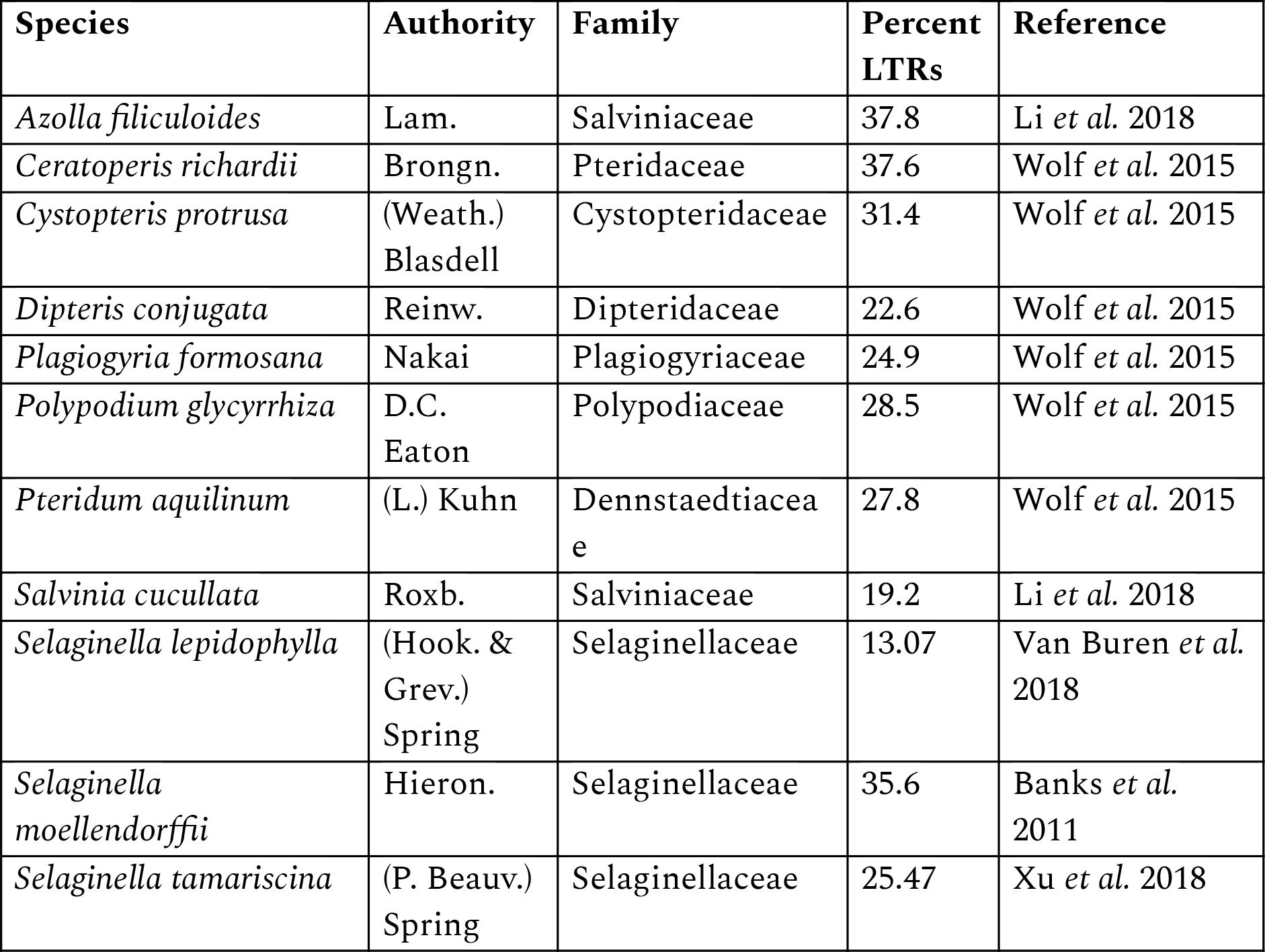
Summary of LTR Composition in Fern and Lycophyte Genomes. Percent of the genome comprised of LTRs was sourced from the literature.

**Table 2.** LTR Summary Statistics. Number of full length LTRs recovered in the genome of each species from our analysis. Arithmetic mean and standard deviation of estimated LTR insertion dates based on lineage specific substitution rates (Appendix 1). Summaries reported for both K2P and GTR + **γ** nucleotide substitution models.

| **Species** | **Authority** | **Haploid**<br>**Nuclear**<br>**Genome Size**<br>**(Mbp)** | **Full**<br>**Length**<br>**LTRs** | **GTR +**<br>**$\gamma$ Mean** | **K2P**<br>**Mean** |
| --- | --- | --- | --- | --- | --- |
| *Selaginella lepidophylla* | (Hook. & Grev.) Spring | 109 | 797 | 0.86 | 1.49 |
| *Selaginella moellendorffii* | Hieron. | 86 | 2,159 | 1.88 | 3.27 |
| *Selaginella tamariscina* | (P. Beauv.) Spring | 150 | 2,675 | 1.73 | 3.04 |
| *Azolla filiculoides* | Lam. | 750 | 17,186 | 1.70 | 3.04 |
| *Ceratopteris richardii* | Brongn. | 11,205 | 125 | 8.18 | 12.58 |
| *Cystopteris protrusa* | (Weath.) Blasdell | 4,230 | 33 | 9.04 | 16.3 |
| *Dipteris conjugata* | Reinw. | 2,450 | 76 | 8.15 | 14.39 |
| *Plagiogyria formosana* | Nakai | 14,810 | 74 | 7.49 | 13.62 |
| *Pteridium aquilinum* | (L.) Kuhn | 15,650 | 391 | 9.52 | 15.99 |
| *Salvinia cucullata* | Roxb. | 260 | 1,711 | 3.11 | 5.52 |
| *Gnetum montanum* | Markgr. | 4,200 | 10,977 | 9.69 | 16.91 |
| *Picea abies* | (L.) H. Karst | 19,570 | 36,868 | 13.15 | 22.54 |
| *Pinus sylvestris* | L. | 22,474 | 89 | 15.74 | 25.48 |
| *Amborella trichopoda* | Baill. | 868 | 1,166 | 4.16 | 6.75 |
| *Arabidopsis thaliana* | (L.) Heynh. | 135 | 286 | 0.75 | 1.31 |
| *Erythranthe guttata* | (Fisch. ex DC.) G.L. Nesom | 362 | 2,535 | 0.45 | 0.79 |
| *Solanum lycopersicum* | L. | 950 | 3,916 | 1.35 | 2.12 |
| *Sorghum bicolor* | (L.) Moench | 697 | 12,980 | 0.64 | 1.13 |

Assembled genomes were scanned using LTR_Finder (Xu and Wang 2007) with the following parameters; “Maximum distance between LTRs=15000”, “Minimum distance between LTRs=1000”, “Maximum LTR length=3500”, “Minimum LTR length=100”, “Length of exact match pairs=15”, “Extension cutoff=0.5”, “Reliable extension=0.5”. Nested LTRs were excluded from all analyses. Predicted full length LTRs were extracted from scaffolds using custom perl scripts (https://bitbucket.org/barkerlab/baniaga_fern_lycophyte_ltrs) and blasted to RepBase (Bao *et al.* 2015) using the tblastx algorithm. The highest scoring hit to the repbase database from each predicted LTR sequence was used in annotations. Predicted full length LTRs with no significant hit to RepBase were blasted to the GenBank nr database using the tblastx algorithm, and significant hits (e-value < 1e^−5^) to protein coding genes were removed. A custom perl script was used to extract 5’ and 3’ LTR pairs from sequences based on their predicted start and end location from the LTR_Finder output (https://bitbucket.org/barkerlab/baniaga_fern_lycophyte_ltrs).

Unlike most investigations of LTR dynamics in vascular plants, we chose to exclude LTRs with a history of intraelement gene conversion. This is because the estimation of LTR insertion age assumes that the 5’ and 3’ ends of an LTR experience mutations independently after insertion. Intraelement gene conversion in LTRs is common in plant genomes (Cossu *et al.* 2017) and acts to homogenize the 5’ and 3’ ends of an LTR. This leads to a reduction in the amount of sequence divergence, and an underestimation of LTR insertion dates. Our exclusion of LTRs with evidence of intraelement gene conversion confronts this potential bias. LTRs with significant evidence of intraelement gene conversion (*p*<0.05) were identified with GENECONV (Sawyer 1989) using the following parameters “/w123/lp/f/eb/g2-include_monosites”. These LTRs were excluded from the dataset and all subsequent analyses.

### LTR insertion date estimation

Insertion dates were calculated following the methods of SanMiguel *et al.* (1998). LTR 5’ and 3’ pairs were aligned in MUSCLE (Edgar 2004) and divergence between LTR pairs was calculated in fastphylo v1.0.1 (Khan *et al.* 2013) using the Kimura two-parameter (K2P) and the GTR+**γ** nucleotide substitution models in FastTree v2.1 (Price *et al.* 2010). Insertion dates were converted into million years using lineage specific synonymous substitution rates per site per year sourced from the literature (Appendix 1), and the arithmetic mean, and standard deviation of insertion dates were calculated for each taxon. In our analyses, the GTR+**γ** nucleotide model consistently estimated younger insertion dates for all taxa, with an average 58% reduction in the mean estimate of LTR insertion date relative to the K2P model. We chose to describe the results of the GTR+**γ** model because it allowed a greater amount of parameters to vary in how nucleotides change over time in comparison to the more simpler K2P model (Tavaré 1986).

### Relationship between mean LTR insertion date and genome size

Haploid nuclear genome size estimates were sourced from the literature. Due to the large variation in haploid genome size across taxa, a regression was performed on mean LTR insertion date against log_10_[haploid nuclear genome size Mbp]. In addition, because of the shared evolutionary history of the species included in our analyses a phylogenetic independent contrast was performed on this dataset. Sequences for the *rbcL* plastid marker were downloaded from NCBI GenBank for each species and the outgroup *Physcomitrella patens* (Appendix 1). An *rbcL* sequence for *Plagiogyria formosana* was not available, and instead a sequence from the congener *P. yakumonticola* Nakaike was used. Sequences were aligned in MUSCLE, and an ultrametric phylogenetic tree was inferred using BEAST v2.0 (Bouckaert *et al.* 2014). The outgroup was pruned from the tree and the phylogenetic independent contrast was performed on both the log_10_[haploid nuclear genome size Mbp] and mean LTR insertion date using the ‘pic’ function in the R ape package (Paradis *et al.* 2004). A nexus file of the phylogenetic tree is available at TreeBASE (http://purl.org/phylo/treebase/phylows/study/TB2:S24023).

## RESULTS

### Fern LTR dynamics

We observed different patterns of LTR dynamics in homosporous versus heterosporous ferns. Across all five homosporous fern taxa analyzed, estimated LTR insertion dates followed a broad unimodal distribution with three common attributes (Fig. 1). Homosporous ferns had few recent LTRs inserted within the past million years. Less than 3% of the LTR insertions occurred within the past one million years in *Ceratopteris richardii* and *Pteridium aquilinum*, and 1% or less of LTR insertions occurred within the past million years in *Dipteris*, *Plagiogyria*, and *Cystopteris*. More recent activity was found in *Cystopteris* with roughly 12% of the insertions occurring between 1-2 mya. The majority of the LTRs in homosporous ferns were estimated to be inserted between 5-13 mya, with a mean of 7.49 mya in *Plagiogyria formosana*, 8.16 mya in *Ceratopteris*, 9.04 mya in *Cystopteris*, 8.15 mya in *Dipteris*, and 9.52 mya in *Pteridium aquilinum*. Finally, we found numerous LTR insertions beyond 15 mya across all homosporous fern taxa, comprising roughly 10% of the total recovered LTR insertions for each taxon. The oldest full length LTR recovered for each taxon was 39.28 mya in *Pteridium*, 34.57 mya in *Ceratopteris*, 28.68 mya in *Dipteris*, 25.42 mya in *Cystopteris* and 24.99 mya in *Plagiogyria*.

**Figure 1.**
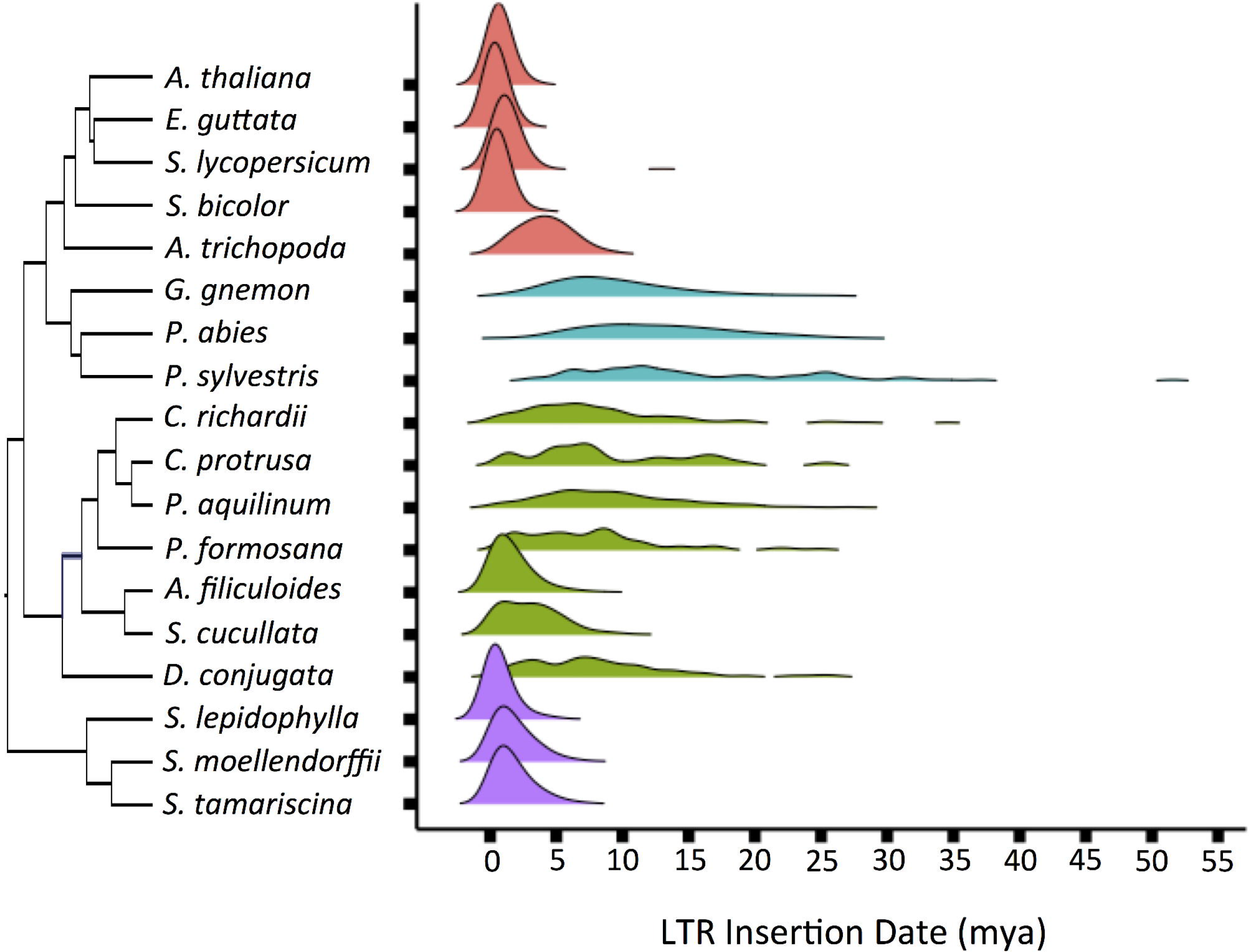
Estimated Distribution of LTR Insertion Times in Vascular Plants. Ridgeline density plots of species across vascular plants with a focus on ferns and lycophytes. The heterosporous *Selaginella* and members of Salviniaceae share similar LTR insertion dynamics with angiosperms. Homosporous ferns share similar dynamics to gymnosperms. Minimum density displayed 0.005, LTR insertion dates are based on the GTR + **γ** nucleotide substitution model and lineage specific substitution rates.

LTR dynamics in the two heterosporous ferns were markedly different from the homosporous ferns. In both *Azolla* and *Salvinia*, estimated LTR insertion dates were similar to those observed in angiosperms with high amounts of recent activity (Fig. 1). In contrast to the homosporous ferns, an estimated 44% of LTR insertions occurred within the past one million years in *Azolla* and 24% in *Salvinia*. Mean insertion dates were also much younger with an estimated mean of 1.70 mya in *Azolla* and 3.11 mya in *Salvinia*. In addition, fewer than 0.1% of LTRs were older than 15 mya in *Azolla* and *Salvinia*.

### Lycophyte (Selaginella) LTR dynamics

All three *Selaginella* species shared similar patterns of LTR activity. These LTR dynamics are comparable to angiosperms and the heterosporous ferns (Fig. 1), in which a large proportion of the estimated LTRs insertions occurred 0-4 mya. Mean LTR insertion dates for the *Selaginella* species were 0.86 mya in *S. lepidophylla,* 1.73 mya in *S. tamariscina,* 1.88 mya in *S. moellendorffii*. In all *Selaginella* genomes sampled, a majority of LTRs have insertion dates within the past 1 million years. An estimated 39% of LTRs are inserted within the past 1 million years in *S. moellendorffii*, 41% in *S. tamariscina,* and 75% in *S. lepidophylla*. Few LTRs inserted beyond 10 mya were recovered for any *Selaginella* species, the oldest estimated LTR insertion for *S. lepidophylla* was 20.75 mya, 22.2 mya for *S. moellendorffii*, and 11.58 mya for *S. tamariscina*.

### Relationship between mean LTR insertion date and genome size

A large range of mean LTR insertion dates and genome sizes were observed in our analyses. Mean LTR insertion dates range from 0.45 mya in *Erythranthe guttata* to 15.74 mya in *Pinus sylvestris*, and genome sizes range from 86 Mbp in *Selaginella moellendorffii* to 22.47 Gbp in *Pinus sylvestris*. A strong significant positive correlation is observed between the haploid nuclear genome size and mean LTR insertion date (Fig. 2). Species with small genome sizes consistently have relatively recent mean LTR insertion dates, and species with large genomes have older mean LTR insertion dates. This is evident for the linear regression of mean LTR insertion date against log_10_[haploid nuclear genome size] (Adj-R^2^=0.77, y=5.012x - 10.3312, *p*<<0.01, df=16, f-statistic=56.89), as well as when phylogenetic relationships are taken into account (Adj-R^2^=0.55, y=4.132x, *p*<<0.01, df=16, f-statistic=21.81).

**Figure 2.**
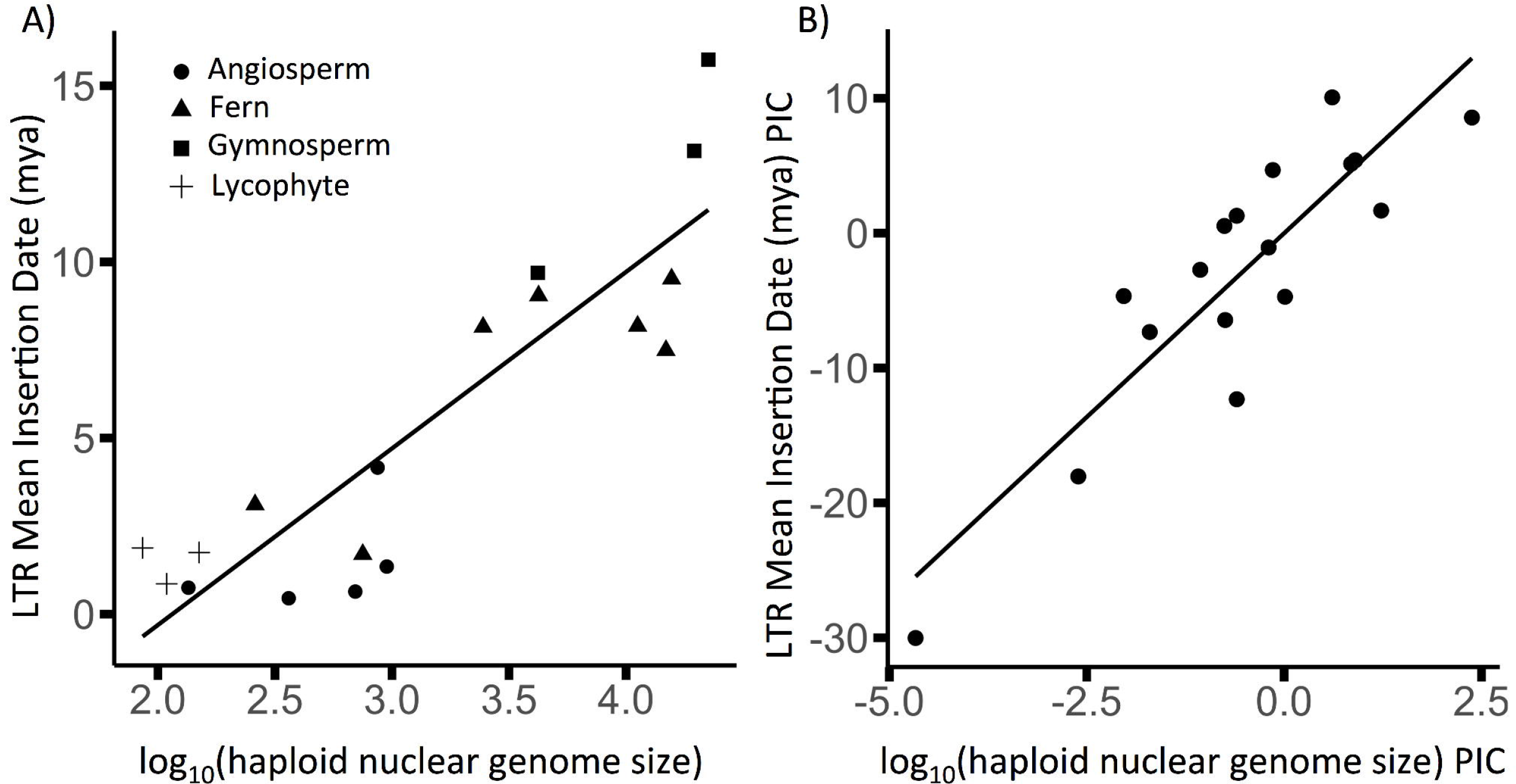
Genome Size by LTR Decay Rate. Species with small genome sizes consistently have relatively recent LTR insertion dates, and species with large genomes have relatively older LTR insertion dates. A) The log_10_(haploid nuclear genome size Mbp) and mean LTR insertion date (mya) across vascular plants. The mean age of LTR insertions is significantly correlated with genome size (Adj-R^2^=0.77, y=5.012x - 10.3312, *p*<<0.01, df=16, f-statistic=56.89). B) The phylogenetically corrected log_10_(haploid nuclear genome size Mbp) and mean LTR insertion date (mya) across vascular plants. The phylogenetically corrected mean age of LTR insertions is also significantly correlated with genome size (Adj-R^2^=0.55, y=4.132x, *p*<<0.01, df=16, f-statistic=21.81).

## DISSUSSION

Across the ten fern and lycophyte genomes we analyzed, we found that haploid nuclear genome size was positively correlated with mean LTR insertion dates. Combined with data from representative seed plants, we found that the age distribution of LTR insertions was well correlated with vascular plant haploid nuclear genome size. Across vascular plants, we found that species with large genomes have older mean LTR insertion dates such as those seen in gymnosperms and homosporous ferns. Species with relatively small genomes had more recently inserted LTRs such as those seen in *Selaginella* and the heterosporous ferns. This first survey of LTR insertion dates for homosporous ferns and other species bridges a significant phylogenetic gap in understanding the evolution of vascular plant genome size. This result is consistent with previous research from seed plants that found genome size and LTR activity are well correlated.

Previous analyses have found a strong positive correlation between haploid nuclear genome size and chromosome number in ferns and lycophytes (Bainard *et al.* 2011; Barker 2013; Clark *et al.* 2016; Leitch and Leitch 2013; Nakazato *et al.* 2008). This pattern is absent from seed plants (Nakazato *et al.* 2008), and instead most of the variation in their nuclear genome size is due to LTR activity (El Baidouri and Panaud 2013; Michael 2014). The high correlation of chromosome number and genome size combined with uniquely small and uniform sized chromosomes in ferns and lycophytes (Nakazato *et al.* 2008; Wagner and Wagner, 1980;) suggested that transposable element activity may be relatively low and not contribute significantly to genome size variation (Nakazato *et al.* 2008). However, our results suggest that LTR dynamics also strongly influence the nuclear genome sizes of ferns and lycophytes (*Selaginella* spp.).

Our current results indicate that the relationship among chromosome number, LTR activity, and genome size is not clearly understood in these plants. It may be the rate of LTR activity is sufficiently low enough that it does not interfere with the strong correlation of genome size and chromosome number generated by the (unknown) forces driving chromosome size uniformity. We currently have too few sampled fern and lycophyte genomes to formally analyze if chromosome number or LTR activity explains more variation in genome size, as well as their joint effect. However, the proportion of variation in genome size explained by LTR activity across the fern and lycophyte species we sampled (adj-R^2^=0.39) is positive but possibly less than that explained by chromosome number (e.g., R^2^=0.83 Nakazato *et al.* 2008; rho=0.5 Clark *et al.* 2016). Future analyses leveraging better sampled fern and lycophyte genomes are needed to explore the relationships among chromosome number, LTR activity, and genome size. Comparative and population genomic approaches of LTR activity and chromosomal evolution are needed in this clade to understand the unique pattern of genome evolution.

Previous analyses have proposed that the high correlation of genome size and chromosome number, as well as the uniform size of fern chromosomes (Nakazato *et al.* 2008; Wagner and Wagner 1980;), is caused by the suppression of transposable elements in homosporous ferns (Nakazato *et al.* 2008). The older average LTR insertion times we observed in homosporous ferns may support this hypothesis. The lifecycle of a LTR inside a host genome involves replication, insertion into a different part of the genome, and potential silencing and deactivation of its activity via epigenetic and methylation processes from the host genome. The position of LTRs inside the host genome can affect the expression of proximal genes, as genes proximal to methylated LTRs often experience higher methylation and reduced expression (Hollister and Gaut 2009; Niederhuth *et al.* 2016). Cytosine methylation is one type of DNA methylation. Across vascular plants, the highest cytosine methylation found to date is in homosporous ferns with an estimated 57% of their genes having greater than 90% of their cytosines methylated (Takuno *et al.* 2016). In contrast, cytosine methylation in *Selaginella* appears to be localized to a subset of approximately 10% of genes (Takuno *et al.* 2016; VanBuren *et al.* 2018). The reason for high methylation in homosporous ferns is unclear. However, homosporous ferns are unique among vascular plants because following whole genome duplication chromosomes appear to be largely retained (Barker and Wolf 2010; Barker 2013; Nakazato *et al.* 2008). It may be that the high methylation in homosporous ferns is associated with silencing LTRs and genes on these retained chromosomes. Future analyses of homosporous fern genomes are needed to disentangle the relationships among these processes and determine the forces ultimately driving their unique patterns of genome evolution.

Older average insertion times of LTRs is consistent with observed patterns of structural turnover in homosporous fern genomes. Unlike many angiosperm lineages, ferns and lycophytes have highly uniform chromosomes in both size and structure (Wagner and Wagner 1980), and some lineages exhibit evidence of conserved genome size over long periods of geologic time (Baniaga *et al.* 2016; Clark *et al.* 2016; Schneider *et al.* 2015). Even fossilized and extant chromosomes of the royal ferns (Osmundaceae) do not show significant difference in size despite more than 180 million years (Bomfleur *et al.* 2014), and sterile hybrids may be generated between species of different genera that diverged more than 60 million years ago (Rothfels *et al.* 2015). Our observation of older average insertion times of LTRs in homosporous ferns is consistent with a slow rate of structural turnover in their genomes.

Why do we observe less active LTRs in homosporous fern genomes? A possible explanation is their unique mating systems. Although most diploid homosporous ferns sampled to date are highly outcrossing (Haufler 1987; Sessa *et al.* 2016; Soltis and Soltis 1992), homosporous ferns are capable of sporophytic and gametophytic selfing (Haufler *et al.* 2016; Klekowski and Baker 1966; Klekowski and Lloyd 1968). Mating system plays a large role in transposable element (TE) dynamics, of which LTRs are one type (Agren *et al.* 2014; Agren and Clark 2018; Boutin *et al.* 2012; Charlesworth and Langley 1986). Theoretical models suggest that if TEs are deleterious to host fitness, the activity and accumulation of TEs should be reduced in inbreeding species. This is because selfers reduce the spread of TEs between individuals, and, compared to outcrossers, more efficiently purge insertions with deleterious recessive effects (Morgan 2001; Wright *et al.* 2008; Wright and Schoen 1999). Outcrossing may also increase the activity and genetic diversity of LTRs during transposition burst cycles (Sanchez *et al.* 2017). Expectations of these models parallel those concerning genetic load in homosporous ferns (Hedrick 1987), and TEs, if deleterious, may comprise a large component of genetic load.

Applying these theoretical models of mating system and TE dynamics to homosporous ferns we would expect that strongly outcrossing species should have more recent LTR insertional activity and species with a history of inbreeding via selfing should have fewer recent LTR insertions. Interestingly, despite relatively limited sampling, our analyses of LTR dynamics in ferns and lycophytes are consistent with these predictions. The heterosporous *Selaginella* and the two aquatic ferns *Azolla filiculoides* and *Salvinia cucullata* share similar LTR dynamics characterized by a predominance of recent insertions. Although there may be several adaptive roles of heterospory in vascular plants (Petersen and Burd 2018), it does function to promote outcrossing. In contrast, LTR dynamics in the homosporous ferns which are capable of a greater array of selfing mechanisms, have relatively fewer recent insertions. As the costs to sequencing decrease it would be worthwhile to experimentally manipulate and compare the TE and LTR dynamics of homosporous ferns with different mating systems such as *C. richardii* and its sister species to empirically verify at a finer scale these macroevolutionary patterns. Similarly, it would also be useful to compare LTR dynamics in homosporous ferns with high rates of gametophytic selfing, such as species in the Ophioglossaceae (Barker and Hauk 2003), to species with low rates of self-fertilization. Ferns and lycophytes may prove to fertile ground for testing hypotheses of LTR dynamics and mating systems in general because of their diverse mating systems.

Another interesting result of our analysis is evidence that we did not observe an effect of generation time on the insertion dynamics of LTRs in homosporous ferns. For example, the minimum generation time of *P. aquilinum* is two to three years (Klekowski 1972), whereas in *C. richardii* the minimum generation time is three months (Banks 1999; Hickok *et al.* 1995; Hickok and Warne 2004). If generation time had a strong effect on activity of LTRs, we would expect to observe a greater amount of recent LTR insertions in *C. richardii*, or a much older mean LTR insertion date in the other fern taxa sampled. Instead, we found that *C. richardii* has a similar LTR insertion distribution as observed in other fern taxa.

LTRs are important components of vascular plant genomes. Our analysis investigated the LTR dynamics of ferns and lycophytes and placed the results in the context of LTR dynamics in other vascular plants. Both the heterosporous *Selaginella* and members of the Salviniaceae share similar LTR insertion patterns to angiosperms, characterized by a predominance of insertions up to 5 mya. In contrast, the homosporous ferns share older average LTR insertion dates intermediate between angiosperms and gymnosperms. We also show that across vascular plants LTR insertion dates are strongly associated with nuclear genome size. Species with small genomes have rapid LTR turnover rates with many young insertions, while species with larger genomes consistently have older LTRs with relatively little recent activity. Ferns and lycophytes are located at key locations to understand LTR dynamics in vascular plants, and elements of their reproductive process such as the capacity for gametophytic selfing in homosporous ferns make them unique study systems for future inquiries into LTR and other transposable element dynamics.

## ACKNOWLEDGEMENTS

The author are grateful for L. Zheng and S. A. Jorgensen for comments on earlier versions of the manuscript. This work was supported in part by NSF grants IOS‐1339156 and EF‐1550838 to M.S.B.

## Data Availability

TreeBase

*.nex

BitBucket data and scripts

**Appendix 1.** List of synonymous substitution rates per site per year used to estimate LTR insertion dates. List of GenBank accession numbers used to infer evolutionary relationships for a phylogenetic independent contrast (* no available sequence for *Plagiogyria formosana* was available and the congener *P. yakumonticola* was used).

| **Species** | **Synonymous Substitution Rate/site/year** | **Reference** | **NCBI GenBank Accession Number** |
| --- | --- | --- | --- |
| *Selaginella lepidophylla* | 6.50E-09 | Gaut *et al.* 1996 | AF419051.1 |
| *Selaginella moellendorffii* | 6.50E-09 | Gaut *et al.* 1996 | KY023084.1 |
| *Selaginella tamariscina* | 6.50E-09 | Gaut *et al.* 1996 | AB574655.1 |
| *Azolla filiculoides* | 4.79E-09 | Barker 2009 | KM360662.1 |
| *Ceratopteris richardii* | 4.79E-09 | Barker 2009 | EU352297.1 |
| *Cystopteris protrusa* | 4.79E-09 | Barker 2009 | JX874051.1 |
| *Dipteris conjugata* | 4.79E-09 | Barker 2009 | EF588692.1 |
| *Plagiogyria formosana* * | 4.79E-09 | Barker 2009 | AB574748.1 |
| *Pteridium aquilinum* | 4.79E-09 | Barker 2009 | AY300097.1 |
| *Salvinia cucullata* | 4.79E-09 | Barker 2009 | U05649.1 |
| *Gnetum montanum* | 2.20E-09 | Nystedt *et al.* 2014 | AY056575.1 |
| *Picea abies* | 2.20E-09 | Nystedt *et al.* 2014 | EU364777.1 |
| *Pinus sylvestris* | 2.20E-09 | Nystedt *et al.* 2014 | AB097775.1 |
| *Amborella trichopoda* | 6.50E-09 | Amborella Genome Project 2013 | L12628.2 |
| *Arabidopsis thaliana* | 1.50E-08 | Koch *et al.* 2000 | KU739560.1 |
| *Erythranthe guttata* | 1.50E-08 | Koch *et al.* 2000 | KF997284.1 |
| *Solanum lycopersicum* | 1.30E-08 | Ma and Bennetzen 2004 | XM_010327541.3 |
| *Sorghum bicolor* | 1.30E-08 | Ma and Bennetzen 2004 | AM849341.1 |
| *Physcomitrella patens* | N/A | N/A | AB013656.2 |

